# Early Vision Shapes Recurrent Processing in the Human Visual Cortex

**DOI:** 10.64898/2026.06.16.731263

**Authors:** Carolin Heitmann, Minye Zhan, Madita Linke, Ramesh Kekunnaya, Rick van Hoof, Rainer Goebel, Brigitte Röder

**Affiliations:** Biological Psychology and Neuropsychology, Faculty of Psychology and Human Movement Science, Universität Hamburg, Hamburg, Germany; Sorbonne Université, Inserm, CNRS, Paris Brain Institute, ICM, Hôpital de la Pitié-Salpêtrière, Paris, France; Jasti V. Ramanamma Children’s Eye Care Center, Child Sight Institute, L. V. Prasad Eye Institute, Hyderabad, India; Department of Cognitive Neuroscience, Faculty of Psychology and Neuroscience, Maastricht University, Maastricht, The Netherlands

**Keywords:** congenital cataract, sight restoration, functional brain development, visual development, recurrent processing, fMRI, occlusion paradigm, connective field modeling, convergence magnitude

## Abstract

Recurrent processing involves feedforward, feedback and lateral connections and is thought to allow efficient visual processing. Anatomical and behavioral studies in humans have suggested that feedback connections mature later in development than feedforward connections and thus were proposed to depend to a larger degree on experience. In order to isolate feedforward from feedback activity and to investigate the role of early visual experience, we assessed seven individuals with reversed congenital cataracts and nine sighted controls using an “occlusion paradigm” with 7T magnetic resonance imaging (Smith & Muckli, 2010): Grayscale images of scenes were presented with the lower right quadrant covered by a white rectangle. We examined whether information about category (beaches, buildings, highways) and individual scenes could be extracted from early visual region vertices (V1 – V3) associated with the occluded quadrant of the visual field, in the absence of bottom-up visual input. This was achieved by decoding individual category or scene context utilizing a linear support vector machine. In addition, bidirectional information flow was assessed using connective field modeling. While both groups showed successful decoding of scene and category from vertices receiving bottom-up visual input, the accuracy was higher in normally sighted individuals than in individuals with reversed congenital cataracts. When bottom-up input was removed, decoding of categories remained successful in both groups, but decoding of individual scenes was only possible in normally sighted control individuals. Connective field modeling results indicated a less precise alignment of feedforward and feedback processing during visual stimulation in individuals with reversed congenital cataracts. These findings suggest that early visual experience is crucial for the refinement of feedback activity which in turn is crucial for well-tuned feedforward processing.

---

Adult visual perception is known to be the result of loops of processing, involving feedforward, feedback, and lateral connections. Insights on their development and maturation have been gained from anatomical (Burkhalter, 1993; Burkhalter et al., 1993) and behavioral studies in human infants (Nakashima et al., 2021), as well as from non-human animal studies (Briggs, 2020). Anatomically, Burkhalter (1993) reported mature feedforward connections from V1 to V2 at four months of age, when feedback connections had not yet reach layer 2/3 (Burkhalter, 1993). Recently, a behavioral study employing a visual backward masking paradigm in infants further supported a delayed maturation of feedback connections (Nakashima et al., 2021). In adults, the perception of a target stimulus is impaired when it is presented with a mask that remains on the screen for longer than the target. These visual backward masking effects have been attributed to recurrent processing in early visual cortex (Di Lollo et al., 2000). Nakashima et al. (2021) adapted the visual backward masking task for infants aged 3 to 8 months by employing a preferential looking paradigm using face stimuli: they expected a face preference only if the face was recognized (i.e., not *masked*). In infants younger than 7 months, a face preference was observed in trials with and without masking. In contrast, infants 7 months and older exhibited a face preference only in the condition without a mask. The authors concluded from this pattern of results that recurrent processing in early visual cortex remains immature in humans until about 6 months of age.

In rodents, layer 1 of V1 has received increasing attention as a candidate location for the integration of bottom up and top down processing streams (De Marco García et al., 2015). More specifically, with development, the prevalence of bottom up input onto L1 interneurons has been reported to decrease in favor of top-down input – a process that appears to be dependent on visual input (Burbridge et al., 2024; Ibrahim et al., 2021). Similarly, in two enucleated monkeys it was found that while the synaptogenesis in striate cortex was unaffected by a lack of visual experience, the typical remodeling of layer IV was disrupted: the relative number of excitatory synapses compared to inhibitory synapses did not decrease as it did in typically developed monkeys between the age of 3 months and 3 years (Bourgeois & Rakic, 1993, 1996).

Since synaptic pruning and cortical thickness changes seem to follow similar time courses (Gilmore et al., 2018; Huttenlocher & Dabholkar, 1997; Sowell et al., 2003), cortical thickness changes in early development have been used as an approximation for structural brain development in humans (Feng et al., 2021; Gilmore et al., 2018; Guerreiro, Erfort, et al., 2015; Hölig et al., 2023). Following this reasoning, the consistent finding of a thicker visual cortex in congenitally blind individuals compared to normally sighted controls (Anurova et al., 2015; Hölig et al., 2023) and compared to late blind individuals (Jiang et al., 2009), may thus reflect the result of an impaired remodeling due to a lack of visual experience in the first years of life (see also Faust et al., 2021). Considering that this step in neural maturation of the early visual cortex is crucial for the establishment of its typical recurrent processing (e.g., Lamme & Roelfsema, 2000), early visual deprivation would be expected to result in impaired visual processing that typically involves top-down mediated tuning of early visual cortex (EVC).

Congenital cataract reversal (CC) provides a rare human model for studying the effects of a transient congenital disruption of visual experience. Despite cataract removal surgery restoring the function of the eye, persisting impairments in multiple aspects of visual processing have provided evidence for early sensitive periods in the emergence of visual cortical processing. Behaviorally, CC individuals have shown stronger impairments in higher order visual functions, such as visual feature binding (Putzar et al., 2007), coherent motion processing (Lewis & Maurer, 2009; Rajendran et al., 2020) and face processing (Gandhi et al., 2017; Le Grand et al., 2001; Putzar et al., 2010) than would have been expected from that populations’ lower visual acuity (Rajendran et al., 2020). Moreover, evidence for tuning of receptive fields in V1 by backward connections from downstream areas (Nurminen et al., 2018; Vangeneugden et al., 2019; Znamenskiy et al., 2024) has suggested that impaired feedback processing in CC individuals might contribute to both lower visual acuities and altered fine-scale retinotopy (Heitmann et al., 2023). Interestingly, CC individuals exhibited increased cortical thickness compared to normally sighted controls, reaching levels indistinguishable from those observed in permanently congenitally blind humans (Feng et al., 2021; Guerreiro, Putzar, et al., 2015; Hölig et al., 2023). These results support the idea of an early sensitive period of the structural remodeling of the EVC which might result in impaired functional tuning of recurrent connectivity.

A relatively higher impairment of higher-level visual processing in CC individuals was further supported by electrophysiological studies: Sourav and colleagues (2018) employed event-related potentials and investigated the first visual cortical response, the so called C1 wave. The polarity reversal of the C1 wave for upper vs. lower visual field stimulation (termed C1 effect) has been associated with retinotopic organization in V1 (Di Russo et al., 2002). A significant C1 effect in the CC group was interpreted as evidence for the existence of a functional basic retinotopic organization in V1. Furthermore, the authors speculated that the broader topography in their CC group might indicate less tuned neural responses (Sourav et al., 2018). Crucially the second ERP, the P1, which has been associated with extrastriate processing (Di Russo et al., 2002), was strongly attenuated in the CC group. Sourav et al. (2018) interpreted their results as supporting early monkey work on visual deprivation suggesting that downstream visual areas are more affected by congenital visual deprivation than the primary visual cortex (Hyvärinen et al., 1981). This ERP study has recently been extended by two studies using steady-state visual evoked potentials (SSVEP) (Pant et al., 2023; Pitchaimuthu et al., 2021). In CC individuals, the second harmonic frequency and the intermodulation frequency responses (elicited by the luminance varying stimuli which were horizontally moving at a lower rate) were attenuated or not significant at all, respectively. Since these two responses have been associated with extrastriate and bidirectional processing, the authors suggested that both extrastriate cortex and bidirectional processing might be impaired (Pitchaimuthu et al., 2021). This idea has recently been confirmed in a modified SSVEP study (Pant et al., 2023): Random luminance changes typically elicit an impulse response in the alpha frequency range (see in Pant et al., 2023). This response is considered to reflect an entrainment of endogenous oscillations which were thought to facilitate stimulus-driven processing. This entrainment response was largely reduced in the CC group providing additional evidence for the idea of impaired top-down processing in the visual hierarchy.

In addition to the electrophysiological evidence suggesting impaired recurrent processing in the visual system of CC individuals, we recently provided evidence for an aberrant visual cortical processing hierarchy in the same individuals as included in the present study using a retinotopic mapping paradigm and functional magnetic resonance imaging (fMRI). More specifically, population receptive fields of CC individuals did not show the typical increase in size up the visual cortical hierarchy (i.e., V1 – V2 – V3; Heitmann et al., 2023) which presumably would allow for spatial integration of information essential for high-level visual processing (Grill-Spector et al., 2017).

To isolate feedback-associated activity, we here employed the occlusion paradigm from Smith and Muckli (2010), in which participants watch various categories of scene images (e.g., houses, highways, cars). As the crucial manipulation, one quadrant of the images was covered by a uniform white occluder, so that any brain activity arising from the occluded part of the image must originate from feedback activity. The contribution of lateral connections for context specific activity in the occluded region of interest (ROI) was reduced by removing vertices activated by a stimulus covering the surround of the occluded quadrant (see Morgan et al., 2019). Following the procedure of previous work, we tested whether scene category or individual scenes could be decoded from vertices representing the occluded quadrant as is typically possible from vertices representing non-occluded quadrants (Morgan et al., 2019; Smith & Muckli, 2010). Such evidence for top-down control of V1 activity was recently confirmed by invasive electrophysiology recordings in monkeys who were exposed to the occlusion paradigm (Papale et al., 2023). V1 neurons whose receptive fields were in the occluded region of the image exhibited weaker activity and their decoding accuracy was overall lower than in neurons with receptive fields in the non-occluded quadrants, as in human fMRI studies.

Based on previous findings of an altered functional organization of the visual cortical hierarchy in CC individuals (Heitmann et al., 2023; Pant et al., 2023; Pitchaimuthu et al., 2021), we expected lower decoding accuracies in the occluded ROI in the CC than the SC group. In non-occluded ROIs, we predicted successful decoding of scenes for both groups, although with lower decoding accuracies in the CC group (see Raczy et al. 2025).

Another means to investigate information flow between cortical regions is through connective field modeling (CF modeling, Haak et al., 2013). By finding the vertices in one area whose time courses best explain the activity of a vertex in another area, this method allows estimating how information is integrated in feedforward and in feedback processing direction. As has been successfully demonstrated in patients with posterior neurodegeneration (de Best et al., 2020), connective field modeling is a promising tool for disentangling altered feedforward vs. feedback processing. We predicted less integration of information in feedforward direction (see Heitman et al., 2023) and less tuned (larger spread) activation in feedback direction in individuals with reversed congenital cataracts.

## Method

### Participants

Throughout the data collection for the present study as part of a larger project (Heitmann et al., 2023; Rączy et al., 2025), we collected data from a total of 24 participants (16 normally sighted controls, 8 congenital cataract reversal individuals). All participants included in the present study were between 18 and 65 years old, right-handed, and had no history of neurological disorders or chronic diseases. To assess the effect of congenital, temporary blindness, we recruited eight individuals with a history of complete, dense and bilateral congenital cataracts (CC) which were surgically removed within the first four years of life.

Participants provided written informed consent for taking part in the study and received a reimbursement for travel costs as well as a small monetary compensation for their participation. Ethical approval had been obtained from the Local Ethics Committees of the Faculty of Psychology and Human Movement, University of Hamburg and of the University of Maastricht. The experiment was conducted according to the principles laid down in the Declaration of Helsinki (2013).

On the day of testing, visual acuity was assessed on the logMAR scale (Logarithm of the Minimum Angle of Resolution; lower values indicate better vision) with the Freiburg Visual Acuity & Contrast Test (FrACT; Bach, 2006). For this measurement, participants wore their typical visual aids (i.e., glasses or contact lenses). For scanning, MR compatible glasses were used. For each CC individual MRI compatible glasses were manufactured according to their prescription.

#### Sample Description for the Occlusion Analysis

Of the eight CC participants, one (CC06, see table 1) was excluded from the analysis of the occlusion data because no brain surface vertices survived the inclusion criteria (see MRI and fMRI processing section). In the remaining seven CC participants (*M*_age_ = 41.43 years, *SD*_age_ = 5.80 years, range 32 – 48 years, 4 female), cataract surgery had been performed at an average age of 20 months (range: 6 to 48 months). From the normally sighted control (SC) individuals, nine were included in the occlusion analysis. The other seven SC individuals were excluded because of either preliminary versions of the experimental paradigm (3), problems during data collection such as excessive sleepiness (1) or errors in the scanning sequence (3). The remaining nine normally sighted control (SC) participants were matched in age and sex distribution (*M*_age_ = 39.67 years, *SD*_age_ = 8.57 years, range 31- 56 years, 6 females).

**Table 1.** Participant characteristics and clinical information of congenital cataract reversal (CC) participants.

| Participant | Age at test (years) | Sex | Family history of Cataract | Age at Surgery (months) | Visual Acuity at test (logMAR) | Self-reported visual impairments |
| --- | --- | --- | --- | --- | --- | --- |
| CC01 | 34 | male | yes | 12 | 0.63 | none reported |
| CC02 | 45 | male | yes | 24 | 0.16 | impaired stereopsis |
| CC03 | 48 | female | yes | 48 | 0.58 | nystagmus, impaired visual field and stereopsis |
| CC04 | 45 | female | yes | 9 | 0.68 | nystagmus, impaired visual field and stereopsis |
| CC05 | 32 | female | no | 24 | 0.18 | none reported |
| CC06* | 46 | female | yes | 24 | 1.27 | nystagmus, impaired visual field and stereopsis |
| CC07 | 46 | female | yes | 18 | 1.16 | nystagmus, impaired visual field and stereopsis |
| CC08 | 40 | male | yes | 6 | 0.56 | nystagmus, impaired visual field and stereopsis |

\* Participant CC06 was excluded from the analysis of the occlusion paradigms but included in connective field modeling of the resting state data

Average visual acuity of the seven CC individuals included in the analysis of the occlusion data was 0.56 logMAR (*SD*_Acuity_ = 0.33 logMAR, see Table 1). Average visual acuity of the SC participants considered for this part of the analysis was −0.03 logMAR (*SD* = 0.22 logMAR, see table 1)

#### Sample Description for Connective Field Modeling

In the connective field modeling analysis, all eight CC individuals (*M*_age_ = 42 years, *Sd*_age_ = 5.63 years, range 32- 48 years, 5 females) and 10 SC individuals (*M*_age_ = 39.90 years, *SD*_age_ = 8.17 years, range 26- 56 years, 7 females) were included.

Average visual acuity of the participants included in the connective field (CF) analysis was 0.65 logMAR (SD = 0.37 logMAR (see Table 1) for the CC group and −0.09 logMAR (*SD* = 0.21 logMAR, range: −0.30 – 0.33 logMAR) for the SC group.

### Stimuli and Design

#### Occlusion paradigm

Participants were presented with three visual scene categories (i.e., beaches, buildings, and highways), consisting of three images each (referred to as e.g., beach-1, beach-2, beach-3, see Figure S1). All images were taken from a previously used stimulus set (Morgan et al., 2019) and were selected for their good discriminability considering the visual impairments expected in the CC participants. The images were in grey scale, matched for global luminance and spanned 13.85 × 10.38° of visual angle. The lower right quadrant of the images was occluded by a white box (approx. 6.92° × 5.19° of visual angle). In addition, three types of contrast-reversing checkerboard mapping stimuli were presented (see Figure 1B): The target checkerboard mapped the inner section of the occluded quadrant, while the surround checkerboard mapped the left and upper 2 degrees of the occluded quadrant. The control checkerboard mapped the remaining three quadrants of the visual field, which were not occluded. At the center of the image, a fixation checkerboard (0.48° × 0.48° of visual angle) was presented which had to be monitored for color change. Participants were instructed to report scene category (i.e., one of beach, buildings, or highway) through button press when seeing a color change in the fixation checkerboard. The scene-button assignment was randomized.

**Figure 1.**
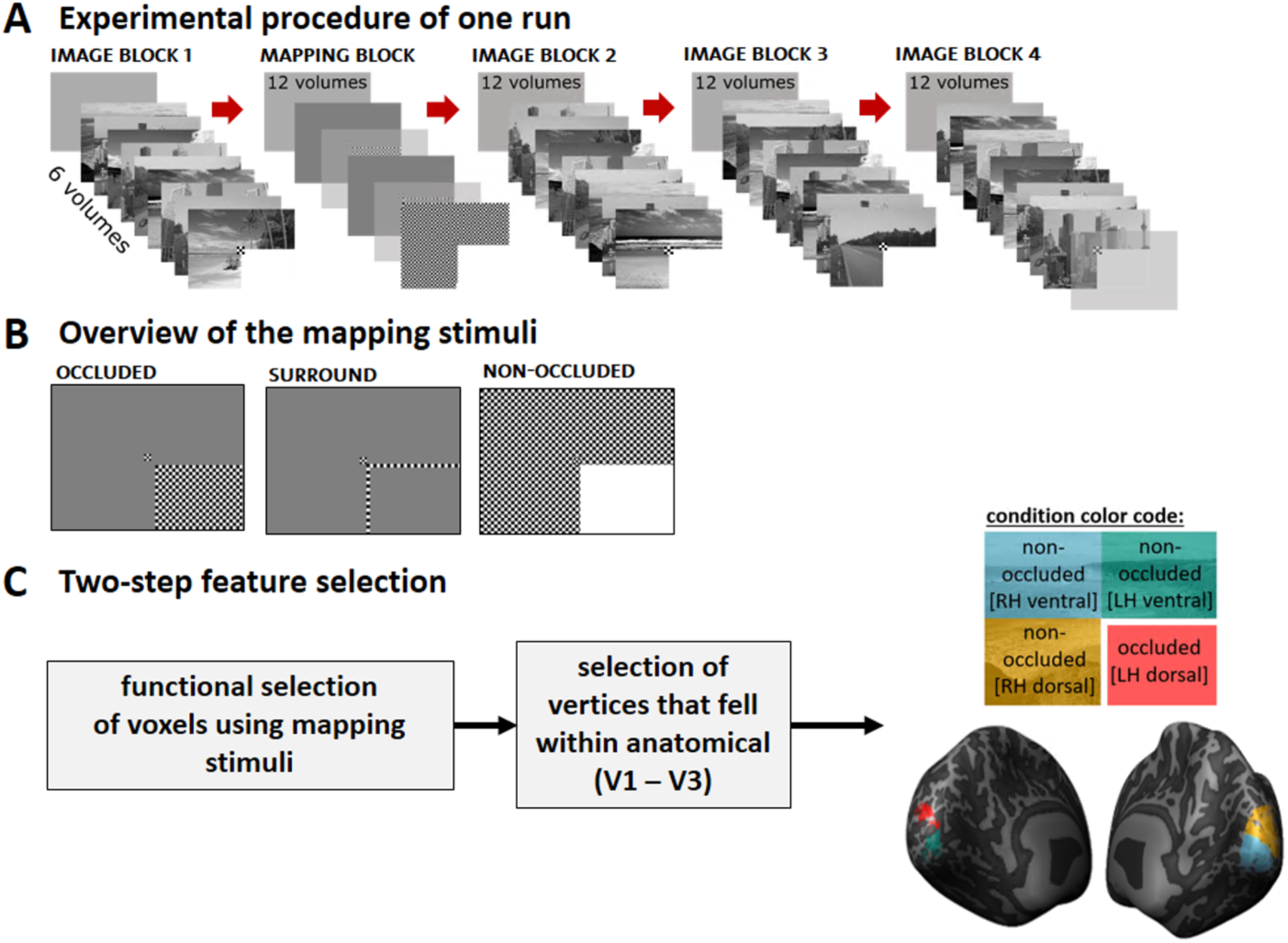
Schematic depiction of the occlusion paradigm and the functional definition of regions of interest ROI). **(A)** Four blocks of images were presented. The first image block was followed by a mapping block in which three retinotopic mapping stimuli were interleaved with baseline periods of no stimulation (gray full-screen). Each image (of the image blocks 1 - 4), mapping stimulus (mapping block) and baseline screen (in gray) was presented for 6 volumes (12 s), unless otherwise specified in the figure. Note that some baseline volumes (i.e. in mapping block and following the last image block) are shown as transparent in this figure to improve visibility. (B) During the mapping block, the three mapping stimuli were presented in fixed sequence: occluded, surround, and control. (C) During feature selection, vertices were first selected based on functional contrasts to determine if they responded to the occluded or to one of the non-occluded quadrants. Next, vertices were restricted to anatomical V1-V3 delineated based on individual participants’ curvature, as well as polar angle data (see Heitmann et al., 2023).

In 12 participants, four runs of 306 volumes (612 s) each were acquired. In the remaining 3 participants (2 SC and 1 CC individual(s)), only three runs were scanned due to time restrictions or participants’ discomfort. Each run included 4 blocks of 9 images (three per category), each presented for 6 volumes (12 s). Images were presented in randomized order within blocks. Each of the blocks lasted for 54 volumes. Since there were four blocks in each run, each run included a total of 216 volumes in which images were presented. In addition, a mapping block was presented\ following the first block of images, resulting in 5 blocks per run (see Figure 1A). During this block, three mapping stimuli (i.e. covering occluded, surround and non-occluded visual field locations as depicted in Figure 1B) were presented for 6 volumes each and interleaved with 6 volumes of baseline. The mapping block thus lasted for 30 volumes. In between blocks, there were 12 volumes (24 s) of baseline. Six volumes of baseline were acquired before the first and after the last block (see Figure 1 for an overview of the paradigm).

#### Population receptive field mapping paradigm

During the population receptive field (pRF) mapping paradigm, participants were asked to look at a red central fixation dot while being presented with a high-contrast bar of checkerboard pattern. The bar was presented in four orientations and jumped randomly between 12 positions per orientation. The bar stimulus covered 10.4° horizontally and vertically. For details on the paradigm, we refer to Heitmann et al., (2023).

### MRI Data Acquisition

Anatomical and functional data were acquired with a 1 transmitter / 32 receiver Nova head coil (Nova Medical, Wilmington, USA) in a 7 Tesla Siemens Magnetom Scanner at Maastricht University (Scannexus, Maastricht, The Netherlands). A high-resolution, whole-brain anatomical scan was collected in a 3D magnetization-prepared 2 rapid gradient echoes (MP2RAGE) sequence in 240 slices (repetition time [TR]: 5000 ms, echo time [TE]: 2.47 ms, voxel size: 0.7mm × 0.7mm × 0.7mm, field of view [FoV]: 224mm × 224mm, flip angle [FA] 1: 5°, FA 2: 3°, inversion time [TI] 1: 900 ms, TI 2: 2750 ms).

Functional occlusion data were acquired with CMRR multiband 2D gradient-echo planar imaging (EPI) sequences in 306 TRs for each run (TR: 2000 ms, TE: 25 ms, voxel size: 0.8 mm × 0.8 mm × 0.8 mm, FoV: 148 mm × 148 mm, FA: 70°, Acquisition matrix: 186 × 186; phase encoding direction anterior to posterior; GRAPPA acceleration factor 3; number of slices: 56, without gaps; Multi-band acceleration factor: 2; echo spacing: 1ms). Slices were oriented to optimally cover the calcarine sulcus and the occipital lobe, which hosts the early visual cortices (V1, V2, V3). Immediately before each functional task run, 5 TRs with identical parameters but opposing phase encoding direction (posterior to anterior) were collected for EPI top-up distortion correction (further described in the next section).

Two runs with 304 TRs (10 min 10 sec) of functional population receptive field data were acquired with the same parameters (see Heitmann et al., 2023 for details).

Resting state data was acquired in one run of 240 TRs (8 minutes) during which the participants were asked to close their eyes and think of nothing in particular.

Besides verbally instructing the participants, foams in the head coil were used to further minimize head movements and increase comfort.

### MRI and fMRI Processing

Structural and functional MRI data were preprocessed and analyzed in BrainVoyager [20.6 and 21.4] (Brain Innovation, Maastricht, The Netherlands) (Goebel et al., 2006), Matlab (version R2021b, The MathWorks Inc., 2021) and Python (version 3.12.3). From the series of 3D volumes acquired with the MP2RAGE sequence, the uniform images were divided by quantitative T1 maps (an optional step to give more similar contrast to the traditional T1 images at lower fields), with the background noise masked out by the inversion 2 images. The resulting anatomical file was adjusted in contrast and brightness, resampled to 0.4 mm × 0.4 mm × 0.4 mm and brought into ACPC space (anterior and posterior commissure). Segmentation of gray matter (GM) - white matter (WM) - cortical spinal fluid (CSF) boundaries was performed with the Advanced Segmentation tools of BrainVoyager, followed by careful manual correction. Next, the GM-WM boundary was morphed into 3D-meshes, on which gyri and sulci were marked in smoothened curvature maps. These meshes were inflated to a sphere, followed by distortion correction vertices to accurately represent curvature of extensively folded regions. Subsequently, each individuals’ spheres (i.e., separately per hemisphere) were sampled onto a high-resolution standard sphere (number of vertices: 163842), and the curvature maps aligned as described in the literature (Frost & Goebel, 2012; Goebel, 2005). Visual areas of interest were taken from our previous study (Heitmann et al., 2023). They had been defined on the basis of calculated individual polar angle maps (i.e., according to meridians indicating visual area boundaries) and eccentricity data (Benson et al., 2014).

Functional data were de-noised using NORDIC (Vizioli et al., 2021) and corrected for EPI distortions by the top-up method in the COPE plugin (Breman et al., 2020) in BrainVoyager, where the voxel displacement maps between the task run and its distortion correction run was estimated and applied (Andersson et al., 2003). Subsequently, functional data were preprocessed in BrainVoyager (slice scan time correction according to the slice timetable in the DICOM headers [sinc interpolation], 3D motion correction [trilinear for estimation/sinc for interpolation], high-pass temporal filtering with Fourier basis set [2 cycles]). No spatial smoothing was applied. Precise manual coregistration to the anatomical data and between functional runs was achieved, by first coregistering one single functional run to the anatomical data, then coregistering all the other functional runs to this run. Volume time courses of functional fMRI data were created in ACPC space with fixed bounding box parameters of the first run. These volume time courses were then sampled across gray matter depths and projected onto the standard high-resolution surface in Brain Voyager (163842 vertices).

Vertices were selected for classifier analysis based on a combination of their functional response during the mapping block and anatomical location (see Figure 1B-C). First, to obtain vertices’ functional responses to different parts of the visual field and thus to differentiate between those responding to the occluded quadrant or to non-occluded quadrants of the visual field, a general linear model (GLM) with predictors for each conditions in the occlusion experiments (all nine scenes [*or* three categories], the three mapping stimuli [see Fig. 1B], and button presses) was fitted to the data across all runs, while regressing out the 6 motion parameters. Vertices were selected as *occluded* vertices if they responded to the occluded quadrant but were not activated by either the non-occluded quadrants of the visual field or by the surrounding area of the occluded quadrant (contrasts: occluded > baseline AND [NOT non-occluded > baseline] AND [NOT surround > baseline], Figure 1B). Vertices were selected as *non-occluded* vertices if they responded to the non-occluded quadrants of the visual field but not to the occluded quadrant or its surrounding area (contrasts: non-occluded > baseline AND [NOT occluded > baseline] AND [NOT surround > baseline], Figure 1B). In addition, early visual cortex ROIs had been defined in our previous study (Heitmann et al., 2023). The definition was based on individual anatomy (as in Benson et al., 2014) and corrected with individual polar angle maps wherever possible and needed. Since we did not expect differences between the three early visual regions V1 to V3 we considered them together in the occlusion analysis to counteract low vertex numbers. Occluded vertices that fell within dorsal left V1-V3 are referred to as *occluded ROI*. Notably, the occluded condition always corresponds to left dorsal V1-V3 since the occluder covered the lower right quadrant of the visual field throughout the experiment. Non-occluded vertices were selected as non-occluded

ROIs, including those within the ventral V1-V3 of the left hemisphere (*non-occluded LV ROI*), dorsal V1-V3 of the right hemisphere (*non-occluded RD ROI*) or right ventral V1-V3 (*non-occluded RV ROI*).

While pRF mapping data were preprocessed as the occlusion data (described Heitmann et al., 2023) additional steps of preprocessing were added for CF modelling of the resting state data. Following recommendations by Power et al. (2014), we regressed out motion parameters (i.e., 3 translational and 3 rotational parameters) as well as average time courses of voxels covering cerebrospinal fluid and those covering white matter. Additionally, to identify and exclude volumes affected by large motion, we applied an upper threshold of 0.3 to individuals’ Framewise Displacement values.

### Multivariate Pattern Analysis

To derive single-trial response estimates for each vertex within each ROI, the GLMsingle package in Matlab (R2021b) was used (Prince et al., 2022). Since hemodynamic response functions may vary due to vasculature and considering the event-related nature of the experimental design, we opted for the “library of HRF” technique, where, for each vertex, a custom hemodynamic response function is automatically selected from a library of candidate functions, according to best fit. Next, a multivariate pattern analysis on the resulting surface maps resulted in a *nr of trials* (144 in participant with 4 runs [108 for participants with 3 runs]) * *nr of vertices* matrix containing the z-scored beta estimates of each voxel’s response to the 144 [108] scene presentations. Using the scikit-learn module (Pedregosa et al., 2011) in Python (version 3.12.3), a linear support vector machine (SVM) was trained separately for all ROIs to map between the multivariate observations of brain activity and the category (i.e., highway, building, or beach) or individual scene (i.e., first image of highway category, second image of highway category, …, third image of beach category) presented. Classification analyses were performed using a leave-one-run-out procedure in which the classifier was trained on *n (number of runs) – 1* run and tested on the left-out run. This procedure was repeated for the *n* runs of the experiment.

### Connective Field Modeling

Connective field (CF) modeling was implemented in Python (version 3.12.3) using the prfpy package (Aqil & Knapen, 2023) and following the framework described by Haak et al. (2013). The method estimates how activity in a target region can be predicted from signals in a source region.

More specifically, for each vertex in the target region, the preprocessed BOLD time series (acquired either during rest or stimulation [pRF paradigm]) were compared with predicted time courses generated by two-dimensional Gaussian connective field models defined on the cortical surface of the source region. Each Gaussian was parameterized by its center location and spread (σ). Distances between vertices of the source region were computed along the cortical surface and predicted time courses were generated as the weighted sum of source vertex time series according to the Gaussian kernel (Haak et al., 2013).

Model parameters were estimated in an initial grid search across candidate CF centers and σ values (0.5 – 15 mm in steps of 0.5 mm), followed by optimization to maximize the variance explained in the observed BOLD time series of the vertex in the target region. For each target vertex, the best-fitting parameters and corresponding explained variances were retained for further analyses.

In the present analysis, we only included models if they explained more than 15% of the variance in the observed time courses (Haak et al., 2013; Maimon-Mor et al., 2024). Each of the early visual regions identified previously through a combination of anatomical and functional features (i.e., V1 - V3; see Heitmann et al., 2023) served as target and source regions. Direction of mapping was indicated as source > target (i.e., V1>V2 for V1 sampling in V2). CF modeling allows investigating the inter-areal sampling spread. While, so far, CF modeling has mostly been applied in the feedforward (FF) processing direction (i.e., V1>V2, V1>V3), we additional included sampling in feedback (FB) processing direction (i.e., V3>V2, V3>V1; as in de Best et al., 2020). To acknowledge the important role of FB connections in visual processing, we examined the effects of a transient phase of congenital blindness on FF and FB sampling as well as their ratio and association with visual acuity.

### Statistical Analysis

In the occlusion paradigm, participants had to respond to a color change in the fixation checkerboard with one of three buttons, depending on which scene category was presented. For each participant, the proportion of correct button presses was calculated and averaged across all runs. We assessed average behavioral accuracies and the range of the averages, separately for SC and CC individuals. One CC participant did not respond to any of the color changes. This subject was excluded from the behavioral analysis and correlations with pRF measures but was included in the occlusion analysis and CF modeling.

To evaluate whether classification was above chance at the group level, average classification accuracies of the participants were compared to chance level. Since the data were not normally distributed, one-sided Wilcoxon Sign-Rank Tests were employed. Since this is a non-parametric test on the median, in the result section we report medians (Mdn) along with interquartile range (IQR). This was done separately for the occluded (left dorsal) ROI and the non-occluded (left ventral, right dorsal, and right ventral) ROIs of SC and CC participants separately for scene and category decoding, resulting in 16 tests. Next, to evaluate the effect of *group* (between-subject: CC and SC groups) and *ROI* (within-subject: occluded [left dorsal], non-occluded left ventral, non-occluded right dorsal and non-occluded right ventral) on classification accuracy, we ran a linear mixed model on *decoding accuracy* separately for scenes and categories. Since the residuals were normally distributed in both, scene and category decoding models, we did not apply a transformation to the data. In both analyses, *participant* was included as random intercept. Model fit was evaluated once for just the model’s fixed effects (marginal R^2^) and once for the entire model, including fixed and random effects (conditional R^2^) (Nakagawa & Schielzeth, 2013). P-values for the fixed effects (i.e., main effect of group, main effect of ROI, interaction effect of group × ROI) were attained by conducting Likelihood Ratio Tests (following Brown, 2021). Post hoc pairwise comparisons were performed on the estimated marginal means and corrected for multiple comparison using the False Discovery Rate (FDR) correction. All results are reported in percentage of correct classifications in the text and displayed in proportion of correct classifications in the figures.

CF modeling was done separately for resting state (rest condition) and pRF run (stimulation condition). For the CF analysis, vertices were included when the best-fitting models explained more than 15% of the variance in the target vertex’ time course. Following the example by de Best et al. (2020), we calculated the difference between the sampling extents (i.e., CF sizes of best-fitting models) of V1 in V2 and V3 (notated as V1>V2 and V1>V3, respectively), as well as of V2 and V3 in V1 (V1<V2 and V1<V3) to derive the difference between feedforward (FF) and feedback (FB) sampling, a measure termed *convergence magnitude*. Larger sampling extents in the FF processing direction are represented by a positive convergence magnitude (i.e., “convergence”), with negative values in turn suggesting larger sampling extents in FB processing direction (i.e. “divergence”; de Best et al., 2020). In order to first indicate the direction of information integration, one-sample Wilcoxon Sign-Rank Test were calculated separately per group, condition (resting state vs. stimulus) and visual region (V2 for V1>V2-V1<V2 and V3 for V1>V3-V1<V3). Resulting p-values were corrected for multiple comparison using the false discovery rate (FDR; Benjamini & Hochberg, 1995). Next, a linear mixed model was constructed explaining *convergence magnitude* from *group*, *condition* (rest vs. stimulation), and *visual region pair (V1-V2 and V1-V2)* with *participant* included as random intercept. Interaction terms were evaluated using likelihood ratio tests comparing nested linear mixed-effects models. Non-significant interactions were removed from the final model, resulting in a model including main effects of *group*, *condition* and *visual region pair*, as well as an interaction of *group* and *condition*. Significant interaction terms were followed up with post hoc tests of estimated marginal means as provided in the emmeans package (Lenth & Piaskowski, 2025). To disentangle the contribution of FF and FB CF sizes to convergence magnitudes, an additional linear mixed model was constructed explaining *CF sizes* from *group*, *condition*, and *processing direction* (FF vs. FB) with *participant* as random intercept. As before, significant interactions were followed up with post hoc tests of estimated marginal means, specifically to assess group differences. In all post hoc tests, resulting p-values were corrected for multiple comparison using the false discovery rate (FDR; Benjamini & Hochberg, 1995). Finally, Spearman rank correlations were run separately for conditions to assess an association of convergence magnitude and visual acuity (in logMAR) across groups, as well as age at surgery in CC individuals.

To evaluate the potential contributions of different additional demographic and behavioral measures with our dependent variables, Spearman’s rank correlations were calculated between decoding accuracies and convergence magnitudes (and CF sizes in FF and FB direction, if motivated by a priori hypotheses) with visual acuity, and age at surgery (in CC individuals). In addition, two retinotopic variables were extracted from Heitmann et al. (2023): (1) population receptive field (pRF) sizes and (2) cortical magnification factor (CMF), both for V1, V2 and V3. These data were either correlated directly with the decoding accuracies, or used to derive other measures, as explained next: V1 pRF size was correlated with decoding accuracies as an estimate of the highest cortical resolution (i.e., smallest pRF sizes in EVC). Other measures derived from pRF size data were the *feedforward integration* (FFI) *effect* and the *eccentricity effect* (*Δs*). To obtain a measure for feedforward integration, we subtracted each individual’s average V1 pRF size from the average pRF size of V3. Next, the slope of pRF sizes (Δs) from central to parafoveal eccentricities was extracted and averaged across V1, V2 and V3 as a measure of the typically observed gradually increasing pRF size with increasing eccentricity (Δs). This typical pattern would result in a positive slope. Finally, CMF was averaged across EVC (i.e., V1, V2 and V3) as a measure of foveal bias. In addition to retinotopic measures which reflect stimulus-based processing only, we also correlated convergence magnitudes during rest and with visual stimulation with decoding accuracies as measures of bidirectional cortical information flow.

To disentangle the effect of retinotopic parameters (i.e., V1 pRF sizes, CMF, the FFI, and the Δs) and convergence magnitudes (i.e., during rest and stimulation) from that of visual acuity, Spearman partial correlation coefficients were calculated with decoding accuracies (separately for categories and scenes) while controlling for the effect of visual acuity (logMAR).

Finally, to assess a potential contribution of head motion during the data acquisition, we correlated participants’ average Framewise Displacement values with their decoding accuracies for scenes and categories. No significant correlations were observed (all *p* ≥ 0.1 and *r* < |0.44|, see Figure S3).

## Results

### Behavioral Results in the Occlusion Paradigm

During the occlusion sessions, participants were asked to monitor a central fixation checkerboard for color change. When the black-and-white checkerboard flashed in red and green, participants were instructed to press one of three buttons, depending on which of the three categories was presented at that time (i.e., beaches, buildings, highways; button assignment was randomized). Data inspection revealed that two SC participants had remembered wrong category-button associations which was corrected by recoding the response options. With this correction, SC participants correctly responded on average inn 99.3% of the fixation color changes (*range*: 98% - 100%). One CC participant did not respond at all. The remaining 6 CC participants performed correct on average in 96.9% of the cases (*range*: 87.42% - 100%). Of note, the non-responding participant was still included in the occlusion analysis but was excluded for the exploratory correlation analyses.

### Category Decoding Results

Given the non-normality of the decoding accuracies, Wilcoxon Sign-Rank Tests were conducted to assess whether decoding of categories was significantly above chance level (33%). Since these tests compare medians against chance level, we report medians (Mdn) and interquartile ranges (IQR, i.e. the difference between the 75^th^ and 25^th^ percentile of the data) in Table 2. The linear support vector machine classifier was able to decode category context at above-chance-level from both the non-occluded and occluded ROIs in both the SC and CC group: non-occluded ROIs for the SC individuals: *right dorsal* (RD): *Z* = 2.67, *p* < 0.01; *right ventral* (RV): *Z* = 2.67, *p* < 0.01; *left ventral* (LV): *Z* = 2.67, *p* < 0.01; *non-occluded ROIs* for the CC group: *RD*: *Z* = 2.37, *p* < 0.01; *RV*: *Z* = 2.37, *p* < 0.01; *LV*: *Z* = 2.37, *p* < 0.01; *occluded ROI* for the SC group: *Z* = 2.67, *p* = 0.002; *occluded ROI* for the CC individuals: *Z* = 2.37, *p* = 0.01) (see Table 2 and Figure 2A).

**Figure 2.**
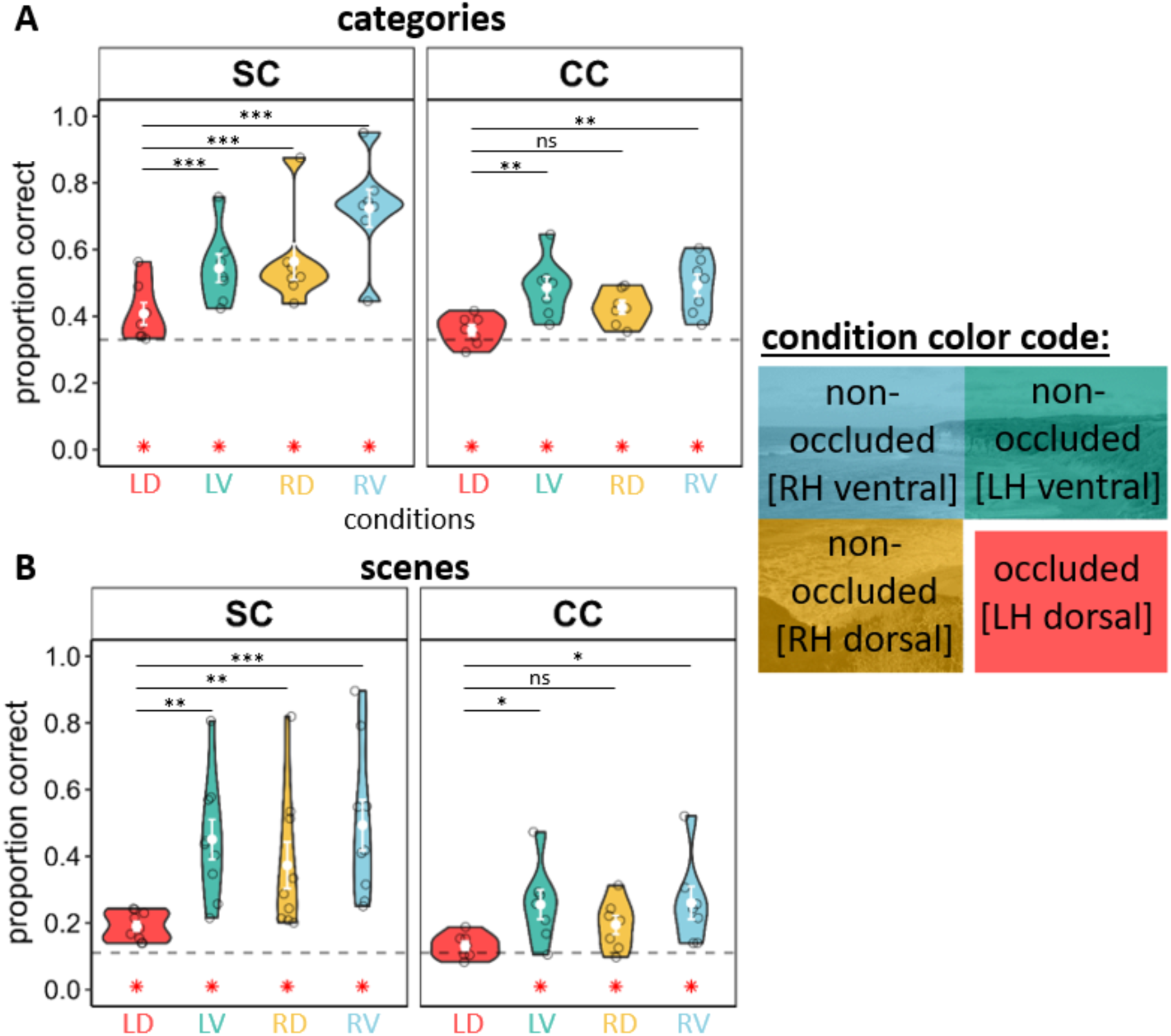
Decoding accuracies per ROI and group. Results of single-trial classification of individual categories (A) and scenes (B) per ROI and group. ROIs are colored according to the colors used for quadrants in the ROI color code subplot on the right. Individual subject, as well as average decoding accuracies are indicated by open circles and filled white dots, respectively. Error bars represent standard errors of the mean. Dashed lines indicate chance level. Red asterisks at y = 0 indicate successful decoding above-chance-level in one-sided Wilcoxon Sign-Rank Tests. Significant pairwise comparisons between ROIs as indicated by paired t-tests corrected for multiple comparison (FDR) are included in black (“ns”: *p* ≥ 0.05, “*”: *p* < 0.05, “**”: *p* < 0.01, “***”: *p* < 0.001).

**Table 2.** Medians and interquartile ranges per ROI and group.

| group | ROIs |  |  |  |
| --- | --- | --- | --- | --- |
|  | occluded<br>(left dorsal, **LD**) | non-occluded<br>(left ventral, **LV**) | non-occluded<br>(right dorsal, **RD**) | non-occluded<br>(right ventral, **RV**) |
| decoding of categories |  |  |  |  |
| normally sighted control individuals | 37.50%**<br>(9.03%) | 51.85%**<br>(6.94%) | 51.85%**<br>(6.94%) | 73.15%**<br>(5.56%) |
| congenital cataract reversal individuals | 36.11%*<br>(5.90%) | 50.00%**<br>(7.99%) | 42.36%**<br>(6.6%) | 51.39%**<br>(12.5%) |
| decoding of scenes |  |  |  |  |
| normally sighted control individuals | 18.52%**<br>(7.64%) | 43.52%**<br>(22.22%) | 28.70%**<br>(30.09%) | 41.67%**<br>(23.38%) |
| congenital cataract reversal individuals | 13.19%<sup>ns</sup><br>(4.86%) | 25.69%*<br>(10.07%) | 20.83%*<br>(9.38%) | 24.31%*<br>(10.42%) |

*Note.* This table depicts medians and interquartile ranges (in parentheses) for decoding of scenes (top part) and categories (bottom part) separately per ROI and group. Differences against chance level (11% and 33% for scenes and categories, respectively) were assessed for statistical significance with Wilcoxon Sign Rank Tests and indicated as follows: “ns”: $p \geq .05$ , “\*”: $p < .05$ , “\*\*”: $p < .01$ , “\*\*\*”: $p < .001$ , “\*\*\*\*”: $p < .0001$ .

To assess the effect of *group* (CC vs. SC; fixed effect, SC as reference) and *ROI* (*occluded LD*, *non-occluded LV*, *non-occluded RD*, and *non-occluded RV*; fixed effect, *occluded LD ROI* as reference) on decoding accuracies, a mixed linear regression model was constructed. Since the residuals of this model did fulfill the normality assumption (*W* = 0.98, *p* = .25), no transformation was applied. Participant was included as random intercept. The fixed effects of the model explained 51.79% (marginal R²; 95% CI [39.24%, 69.20%]) of the variance, which increased to 74.09% with the inclusion of the random intercept (conditional R^2^; 95% CI [62.03%, 84.54%]).

Using likelihood ratio tests comparing nested models, a significant interaction between group and ROI was identified, χ²(3) = 12.46, *p* < .01. Although the main effects of group and ROI were significant as well (both *p* < .01), they were not interpreted due to the presence of the significant interaction. Instead, post-hoc tests on estimated marginal means were conducted. In the SC group decoding of categories was significantly worse in occluded (*M* = 40.48%, *SD* = 7.84%) than in all non-occluded ROIs (all *M* > 54.39%, all *p*_corr_ < .001; see Figure 2A, left panel). A similar pattern of higher decoding accuracies in non-occluded ROIs was found in ventral ROIs (*LV* and *RV* both *M* > 48.61%, all *p*_corr_ < .01; *RD* ROI: *M* = 42.66%, *SD* = 5.18%, *p*_corr_ = .15) of CC individuals compared to the occluded ROI (*M* = 35.81%, *SD* = 4.39%; see Figure 2A, right panel). Comparing the two groups in decoding accuracy in each ROI suggested indistinguishably decoding accuracy in both groups for occluded and one non-occluded ROI (*occluded LD*: *p*_corr_ = .40; *non-occluded LV*: *p*_corr_ = .31), but worse decoding of categories from the two non-occluded CC individuals’ right hemisphere ROIs (*RD*: *p*_corr_ = .03; *RV*: *p*_corr_ < .001).

### Individual Scene Classification Results

As for categories, decoding of scenes was compared against chance level (11%) using Wilcoxon Sign-Rank Tests. Medians and interquartile ranges are reported in Table 2. The linear support vector machine classifier successfully decoded individual scene context from the occluded (*Z* = 2.67, *p* < .01), as well as from all non-occluded ROIs in SC individuals. The highest accuracy was found for the non-occluded left ventral (*Z* = 2.67, *p* < .01), the non-occluded right ventral ROI (*Z* = 2.37, *p* < .01) and the non-occluded right dorsal ROI (*Z* = 2.37, *p* < .01). In CC individuals, decoding was successful only in non-occluded ROIs (all *Z*s > 2.10, all *p*s < .05). In contrast to the SC group, decoding of individual scene context in the occluded ROI was not significantly above chance level (*Z* = 1.18, *p* = .15) (see Table 2 and Figure 2B).

As for categories, *group* and *ROI* were included in a mixed linear regression model with participant as random intercept. The model’s fixed effects explained 41.16% of the variance in decoding accuracy (marginal R²; 95% CI [28.42%, 65.45%]), which increased to 74.79% with the inclusion of the random intercept (conditional R²; 95% CI [61.75%, 85.52%]).

Across ROIs, decoding of individual scene context was significantly higher in SC individuals (M = 37.66%, SD = 20.90%) than in CC individuals (*M* = 21.01%, SD = 10.58%), χ^2^ (1) = 6.56, *p* = .01. Furthermore, decoding accuracies differed between ROIs when assessed across groups (χ^2^ (3) = 34.30, *p* < .001), with the highest accuracy in *non-occluded RV ROI* (*M* = 39.12%, *SD* = 22.06%), followed by *non-occluded LV ROI* (*M* = 36.49%, *SD* = 18.16%) and *RD ROI* (*M* = 29.47%, *SD* = 18.47%). Decoding in *non-occluded RV* was significantly higher than in *RD* (*p*_corr_ = .02). The lowest accuracy was achieved in the *occluded LD ROI* (*M* = 16.42%, *SD* = 4.92%), which was significantly lower than in all *non-occluded ROIs* (all *p*_corr_ < .01). Since the interaction of *group* and *ROI* contributed marginally to the model fit (χ^2^ (3) = 7.29, *p* = .06), post-hoc tests on estimated marginal means were conducted to explicitly assess a priori hypotheses. In SC individuals, the decoding accuracy in the *occluded LD ROI* was 19.01% (*SD* = 4.29%), which was significantly lower than the accuracies in all non-occluded ROIs (*RV*: 49.33%, *SD* = 22.74%, *p*_corr_ < .001; *LV:* 45.04%, *SD* = 18.10%, *p*_corr_ < .001; *RD:* 37.27%, *SD* = 21.02%, *p*_corr_ = .002), see Figure 2B (left panel). In CC individuals, the average decoding accuracy in the *occluded LD ROI* was 13.10% (*SD* = 3.62%), which was significantly lower than the accuracy in the *non-occluded LV* (25.50%, *SD* = 11.70%, *p*_corr_ = .04) and *RV* (26.00%, *SD* = 13.02%, *p*_corr_ = .04) ROIs, see Figure 2B (right panel). The groups did not significantly differ in their decoding accuracies in the occluded ROI (*p*_corr_ = .48), but accuracies were lower in CC individuals in all non-occluded ROIs (all *p*_corr_ < .05).

### Association of Decoding Accuracy with Visual Acuity and Age at Surgery

We calculated Spearman Rank Correlations to assess the associations of decoding accuracies and visual acuity across groups. In addition, we display group-wise correlations in Figure 3. Across groups, better visual acuity was associated with higher decoding accuracies for categories and scenes in all ROIs except for decoding of categories from *occluded LD ROI*. In SC individuals only decoding results from right hemisphere non-occluded ROIs were significantly associated with visual acuity in category decoding. In CC individuals, better visual acuity (i.e., a lower logMAR value) was significantly associated with better decoding of individual scenes in ventral non-occluded ROIs (both *r*s < −0.78, *p* < .05), and with decoding of categories in all non-occluded ROIs (all *r*s < −0.8, *p* < .05).

**Figure 3.**
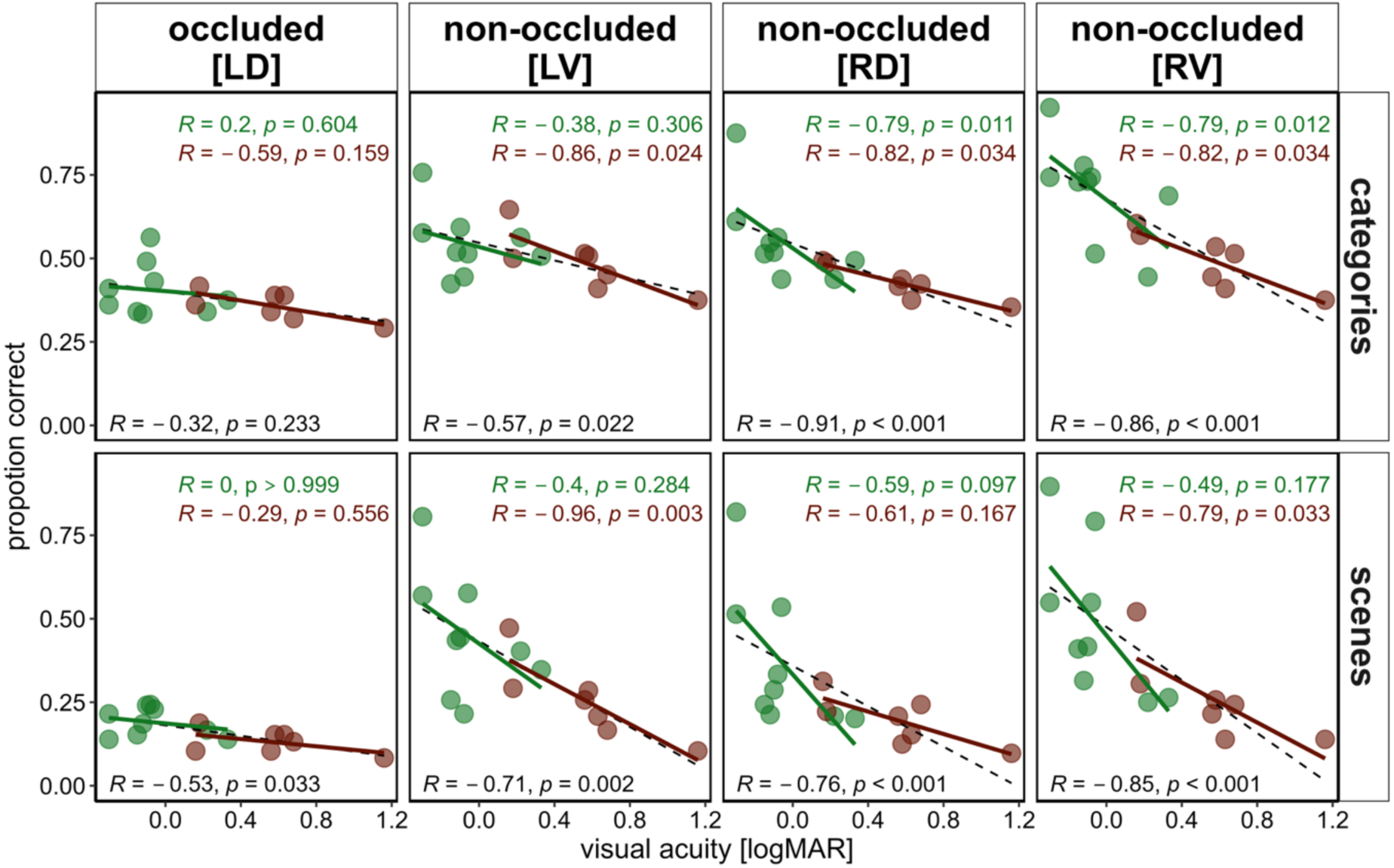
Association of decoding accuracy and visual acuity separately per ROI. Circles represent individual decoding accuracies in proportion of correct classifications for scenes (A) or categories (B) for occluded (left dorsal [LD]) and non-occluded (left ventral [LV] and right dorsal [RD]) regions. Spearman rank correlation coefficients and linear regression lines are displayed separately per group (normally sighted control = green, congenital cataract group = brown), as well as across-groups (black).

Within the CC group, age at surgery was not significantly associated with decoding accuracy (all *p* > 0.17, see Figure S2).

### Connective field modeling results

Convergence magnitude, a measure of the ratio of bidirectional information flow introduced by de Best et al. (2020) and previously described by Haak et al. (2013), was derived by subtracting average connective field (CF) sizes in feedback (FB) direction (i.e., V1<V2, V1<V3) from those in feedforward (FF) direction (i.e., V1>V2 and V1>V3). This was done for CF sizes obtained during visual stimulus processing and during rest with eyes closed. Negative convergence magnitude indicates larger CF sizes in FB direction and hence *divergence* from V1 to V2 or V3 while positive convergence magnitude reflects *convergence* from V1 to V2 or V3. During rest, average convergence magnitude was negative in both groups and for both V2 and V3, suggesting divergent processing with larger CF sizes in FB processing direction in the absence of stimulation (i.e., V1<V2 and V1<V3 larger than V1>V2 and V1>V3; all *Mdn* < −0.34 mm, IQR > 0.28 mm, *p_corr_* < .05, *Z* > 2.16, *r* > .76), see Figure 4A (left panel). During visual stimulation, SC individuals’ convergence magnitude was positive, suggesting larger CF sizes in FF than FB processing direction (V1>V3: *Mdn* = 0.85 mm, *IQR* = 0.62 mm, *p_corr_* = .03, *Z* = 2.34, *r* = .74; not significant: V1>V2: *Mdn* = 0.36 mm, *IQR* = 0.32 mm, *p_corr_* = .13, *Z* = 1.53, *r* = .48). In contrast, CC individuals continued to show negative convergence magnitudes during visual stimulation (V1>V2: *Mdn* = − 0.40 mm, *IQR* = 1.11 mm, *p_corr_* = .03, *Z* = 2.45, *r* = .87; not significant: V1>V3: *Mdn* = −0.28 mm, IQR = 0.91 mm, *p_corr_* = .09, *Z* = 1.75, *r* = .62), see Figure 4A (right panel).

**Figure 4.**
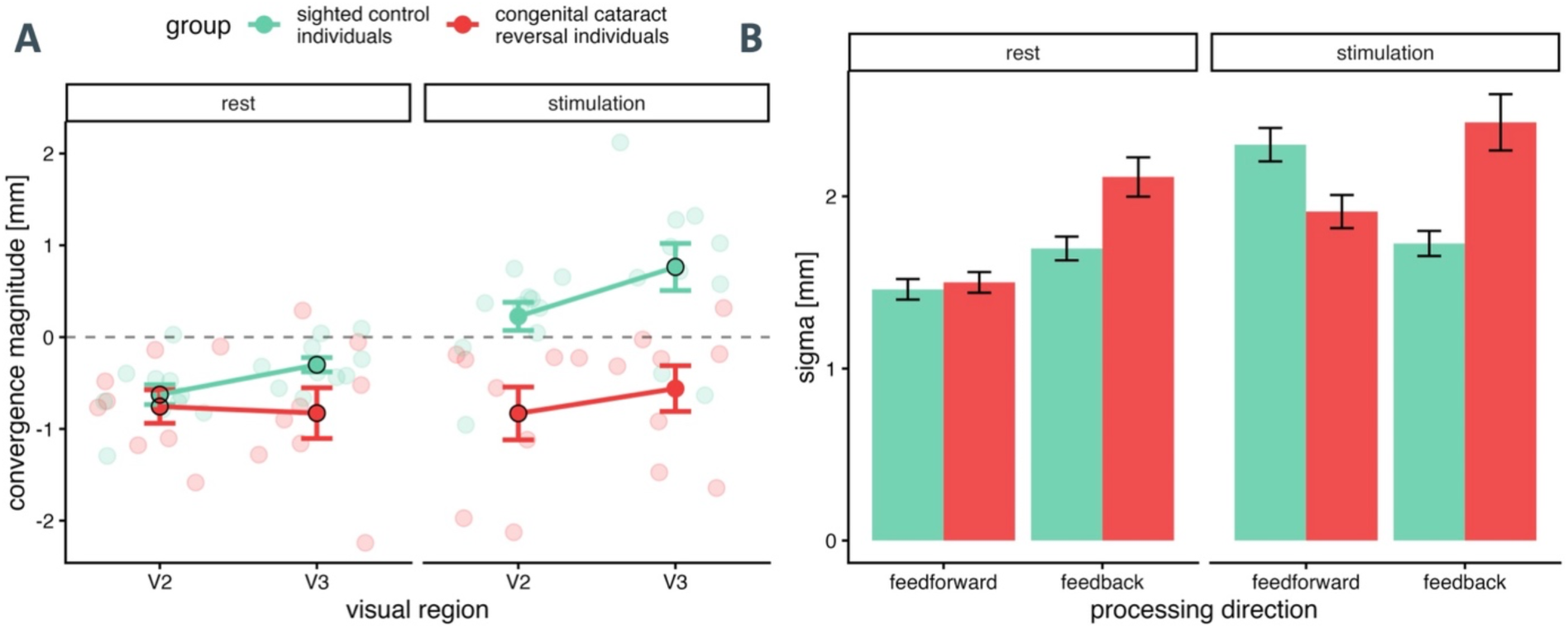
Convergence magnitude from V1 along the visual cortical hierarchy and CF sizes in feedforward and feedback direction per group and condition. (A) Circles indicate mean convergence magnitudes (average feedback [FB] CF size subtracted from average feedforward [FF] CF size; in mm) per group and condition with transparent circles representing individual participant data. Significant one-sample Wilcoxon Sign-Rank Test results against 0 are indicated with black frames (FDR corrected). The horizontal dashed line highlights zero convergence magnitude. (B) Average FF and FB CF sizes (in mm) per group and condition. Error bars represent standard errors of the mean. The sighted control and congenital cataract reversal groups are represented in green and red, respectively.

To evaluate the effect of *group* (SC vs. CC, SC as reference level) and *condition* (rest vs. visual stimulation, rest as reference level) on convergence magnitude, a linear mixed model was constructed with main effects and interactions of these variables. *Visual region pair* (V1V2 vs. V1V3; V1V2 as reference level) was included as main effect only, after likelihood ratio tests confirmed no significant interaction terms (all χ^2^(1) < 1.57, *p* > .21) but a marginally significant main effect (χ^2^(1) = 3.65, *p* = .06) indicating an increase of CF size with hierarchical distance of the visual region pairs (i.e., V1V3 > V1V2; effect in final model: *estimate* = 0.28, *SE* = 0.14, *t*(51) = 2.06, *p* = 0.04). Participant was included as random intercept. The fixed effects of the model explained 44.37% of the variance in convergence magnitude (marginal R^2^; 95% CI [30.52%, 61.84%] which increased to 48.30% with the inclusion of the random intercept (conditional R^²^; 95 % CI [33.69%, 70.16%]). The model suggested a significant interaction of group and condition (χ^2^(1) = 9.72, *p* < .01) which was followed up by pairwise post-hoc tests on estimated marginal means. While mean convergence magnitude was not different in both groups during rest (contrast SC – CC: *estimate* = 0.33 mm, *SE* = .21 mm, *t*(42.20) = 1.56, *p*_corr_ = .12), convergence magnitude increased significantly with visual stimulation in the SC group (contrast of rest – stimulation: *estimate* = −0.96 mm, *SE* = 0.19 mm, *t*(51) = −5.16, *p*_corr_ < .001), but not in the CC group (contrast of rest – stimulation: *estimate* = −0.10 mm, *SE* = 0.21 mm, *t*(51) = −0.47, *p*_corr_ = .64), resulting in significantly larger convergence magnitude of SC than CC individuals during visual stimulation (contrast SC – CC: *estimate* = 1.19 mm, *SE* = .21 mm, *t*(42.2) = 5.64, *p*_corr_ < .001).

To evaluate the contribution of feedforward and feedback CF sizes on the observed results, an additional linear mixed model was constructed to explain the variance in CF sizes with main effects and interactions of *group* (SC vs. CC, SC as reference level), *condition* (rest vs. visual stimulation, rest as reference level) and *processing direction* (FF vs. FB; FF as reference level). Due to the non-normality of this model’s residuals (W = 0.95, p < .01), CF sizes were log transformed (W = 0.99, p = .72). The fixed effects of the resulting model explained 21.49% of the variance in connective field size (marginal R^2^; 95% CI [15.64%, 34.28%] which increased to 38.76% with the inclusion of the random intercept (conditional R^²^; 95 % CI [22.50%, 49.78%]). After the confirmation of a significant three-way interaction between all independent variables of the model through likelihood ratio test (χ^2^(1) = 8.95, *p* < .01), post-hoc paired simple contrasts of *group* on estimated marginal means were conducted while keeping the other variables constant. During rest, CF sizes in the FB direction were larger in CC individuals than in SC individuals (SC – CC: *estimate* = −0.37 mm, *SE* = 0.17 mm, *t*(29) = −2.14, *p*_corr_ = .04, see Figure 4B). During stimulation, CF sizes of CC individuals were significantly larger in FB direction (SC – CC: *estimate* = −0.60 mm, *SE* = 0.19 mm, *t*(29) = −3.19, *p*_corr_ < .01), but marginally smaller than those of SC individuals in FF direction (SC – CC: *estimate* = 0.37 mm, *SE* = 0.19 mm, *t*(29) = 1.97, *p*_corr_ =.06; see Figure 4B).

Spearman rank correlations of convergence magnitude (averaged across visual region pairs) with visual acuity (in logMAR) across groups suggested an association of better visual acuity with higher convergence magnitudes during visual stimulation, but not during rest (see Figure S4A). To disentangle the effect of FF and FB processing in this association, we correlated visual acuity separately with average FF and FB CF sizes: Across groups, better visual acuity was significantly associated with larger CF sizes in FF processing direction and marginally associated with smaller CF sizes in FB direction during visual stimulation (see Figure S4B).

### Exploratory Analysis: Association of Decoding Accuracy, Retinotopic Measures, as well as Convergence Magnitude

Figure 6A shows an overview of Spearman’s Rank correlation coefficients between decoding accuracies for both categories and individual scenes with retinotopic (pRF size- and CMF-derived) measures as well as convergence magnitude during rest and visual stimulation. To reduce complexity and the number of tests, the decoding accuracies of the three *non-occluded ROIs* (*LV*, *RD*, *RV*) were averaged to obtain two conditions: *occluded* and *non-occluded*. Retinotopic measures included were average pRF size in V1 (s [V1]), the slope of pRF sizes from foveal to parafoveal eccentricities (Δs), the increase of average pRF sizes from V1 to V3 (feedforward integration [FFI] effect), as well as average CMF across V1, V2 and V3. These measures quantify cortical resolution (s [V1]), the typical foveal bias (Δs, CMF), as well as spatial integration downstream the visual processing pathway (FFI effect). In addition, convergence magnitude captures the ratio of FF and FB sampling in early visual cortex during rest (rest CM and stimulation (stim. CM).

**Figure 6.**
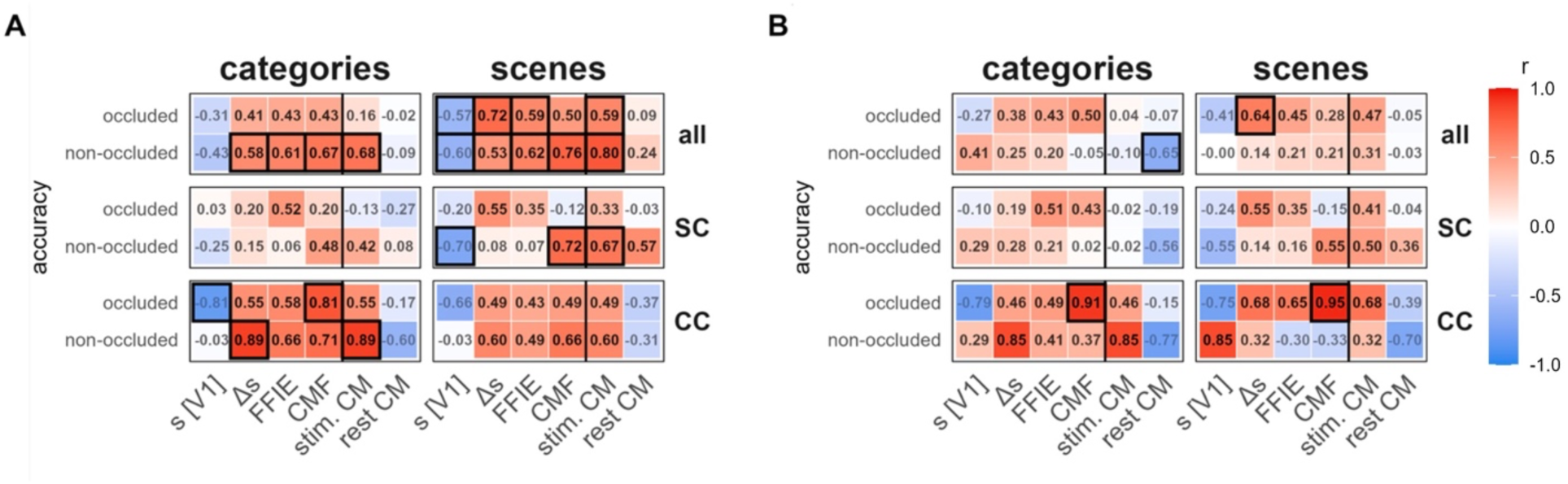
Overview of Spearman’s rank correlation and partial correlation coefficients between decoding accuracies, retinotopic measures and convergence magnitudes per condition across and separately per groups. (A) Spearman rank correlation coefficients between decoding accuracies (i.e., proportion of correct classifications) for categories (left) and individual scenes (right) with average pRF size in V1 (s[V1]), average slope of pRF size with eccentricity across V1 - V3 (Δs; derived by *s ∼ eccentricity*), the feedforward integration effect (FFI effect; the increase of s from V1 to V3), average cortical magnification factor across V1-V3 (CMF), as well as convergence magnitude during rest (rest CM) and stimulation (stim. CM). Correlations are calculated across groups (upper panel), as well as separately per group (middle panel: normally sighted control [SC] individuals; lower panel: congenital cataract reversal individuals [CC]). (B) Partial spearman rank correlation coefficients describing the association of decoding accuracies with the same measures while controlling for visual acuity. Coefficients with associated uncorrected p-values < 0.05 are framed. Note, one CC individual was excluded from this analysis due to highly influential retinotopic measures.

Across groups and for both conditions and levels of decoding (i.e., category or scene), the directions of associations were the same: average pRF size in V1 was negatively associated with decoding accuracy, while pRF size slope (Δs), FFI effect, CMF and stim. CM showed positive associations. All associations except for rest CM reached significance in scene decoding; for category decoding significant positive associations were observed for the *non-occluded* condition. In CC individuals, significant associations were observed in category decoding: smaller V1 pRF sizes and larger average CMF predicted better decoding in the *occluded* condition as well as the slope of pRF size and stim. CM in the *non-occluded* condition. In SC individuals, significant associations were observed in the non-occluded condition of scene decoding: smaller V1 pRF sizes as well as larger CMF and stim. CM predicted better decoding.

As suggested by the associations between visual acuity and retinotopic measures (i.e., pRF size and CMF; see Duncan & Boynton, 2003; Heitmann et al., 2023; Silva et al., 2021) and convergence magnitude, cortical factors majorly contribute to visual acuity. To assess the above-described associations after removing their contribution to visual acuity, we calculated partial spearman rank correlation coefficients controlling for visual acuity (Figure 6B). Across groups, smaller (i.e., more negative) rest CM predicted better decoding of categories in the non-occluded condition while decoding of scenes in the occluded condition was the higher the more positive the slope of pRF size from foveal to parafoveal eccentricities (Δs). In CC individuals, scene and category decoding in the occluded ROI were strongly associated with a larger average CMF (Figure 6B).

We hypothesized that worse decoding of both categories and scenes in non-occluded ROIs of CC individuals might be related to coarse feedback processing. In fact, across groups, smaller FB CF sizes were correlated with better decoding of categories (*r* = −.53, *p* = .05) and scenes (*r* =-.69, *p* < .01) while a similar association was not significant for FF CF sizes (both *r* < .30, *p*s > .28).

## Discussion

Visual perception depends on recurrent processing in the early visual cortex, that is, feedforward, feedback and lateral connectivity. Due to the longer maturation of feedback than feedforward connections (Burkhalter, 1993; Nakashima et al., 2021), we hypothesized that feedback processing particularly depends on early visual experience. Indeed, non-human animal studies have suggested that visual experience in the first months of life is critical for the remodeling of V1 input from a predominantly bottom-up to an orchestrated bottom-up and top-down connectivity (Burbridge et al., 2024; Ibrahim et al., 2021). To directly investigate the influence of early visual experience on feedback processing in humans, seven individuals born with dense bilateral congenital cataracts (CC), whose cataracts had been surgically removed between 6 and 48 months of age, and nine matched normally sighted control (SC) participants were assessed using an occlusion paradigm (Smith & Muckli, 2010). In a high-resolution 7T functional magnetic resonance (fMRI) study, grayscale scenes were presented, with the lower right quadrant blocked by a uniform white occluder to prevent any bottom-up visual input to vertices with receptive fields at these visual field locations. Considering the crucial role of feedback processing in typical visual perception (Briggs, 2020), we expected specific information of categories (i.e., buildings, highways, beaches) and individual scenes within these categories - not only in non-occluded ROIs but additionally in the occluded ROI (as in Morgan et al., 2019; Smith & Muckli, 2010). Because no bottom-up input reached this region and lateral connections are short and cannot explain the effects (see below), any context-specific information is assumed to stem from top-down activation. Considering repeatedly observed lower performance of CC individuals in higher-level visual tasks (Le Grand et al., 2001; Röder & Kekunnaya, 2021), we expected chance-level decoding accuracies in occluded ROIs but an above chance level read-out of category- and scene-specific information in non-occluded ROIs in CC individuals (see Raczy et al., 2025). Note, that most previous neuroscientific or behavioral studies on CC individuals which reported impairments in higher order visual processing did not aim for dissociating the contribution of feedforward and feedback processing (Grady et al., 2014; Heitmann et al., 2023; Ossandón et al., 2025; Rączy et al., 2025; Sourav et al., 2018).

We replicated results of previous studies demonstrating above-chance level decoding of categories and individual scenes in both non-occluded and occluded ROIs in SC individuals, suggesting that occluded, non-stimulated vertices contained detailed information specific to the surrounding visual scene information (Morgan et al., 2019; Smith & Muckli, 2010). In contrast, CC participants showed successful decoding from non-occluded ROIs in scene and category decoding, while only categories, but not individual scenes, could be decoded above-chance level in the occluded ROI. Moreover, CC individuals had overall lower decoding accuracies of both scenes and categories across occluded and non-occluded ROIs compared to SC individuals. These differences were significant in non-occluded ROIs. Across groups, decoding from non-occluded ROIs was significantly higher than decoding from occluded ROIs. Exploratory correlation analyses after controlling for visual acuity indicated a positive association of decoding accuracy and the cortical magnification factor in the CC group.

Connective field modeling results replicated a state dependent shift in processing from divergent processing (i.e., larger feedback than feedforward CF sizes) during rest to convergent processing (i.e., larger feedforward than feedback CF sizes) during visual stimulation in early visual cortex of SC individuals (de Best et al., 2020). In contrast, CC individuals exhibited a “divergent state” independent of whether participants were exposed to visual stimulation. Across groups, this ratio of FF and FB CF sizes during visual stimulation predicted both visual acuity as well as decoding accuracy in the occlusion paradigm.

In sum, two independent approaches to assess the dependence of feedback processing and the interaction of feedback and feedforward processing on early visual experience provided evidence for a disruption of feedback processing with direct consequences for feedforward processing, as discussed below.

### After a transient phase of congenital blindness feedback processing in visual cortex is coarser

Above-chance decoding of categories and individual scenes from responses in neural populations associated with occluded parts of the visual field suggests that these vertices hold information on the surrounding context. Given that feedforward information is obstructed in these neural populations, context-specific information must be assumed to stem from feedback (top-down) connections (Smith & Muckli, 2010). Notably, feedforward and feedback connections link neuronal populations across the visual processing hierarchy and process overlapping parts of the visual field. The influence of lateral (horizontal) connections, which enable interactions between neural units which process non-overlapping parts of the visual field (Angelucci & Bressloff, 2006; Lamme & Roelfsema, 2000) is deliberately reduced in the occlusion paradigm by eliminating vertices responding to the immediate surround of the occluded part as well as the non-occluded parts of the images (see Figure 1B). Consequently, according to Smith and Muckli (2010), information present in the finally selected vertices of the occluded ROI must predominantly stem from feedback activity.

The lack of above-chance individual scene decoding in occluded ROIs for CC individuals hence suggests impaired feedback processing, as we had predicted based on previous behavioral (Le Grand et al., 2001; McKyton et al., 2015; Putzar et al., 2007) and electrophysiological results (Pant et al., 2023; Pitchaimuthu et al., 2021). However, the successful decoding of category context in the same individuals indicated some level of functional feedback processing in CC individuals. A previous fMRI study in CC individuals had demonstrated that area MT exhibited higher levels of activity when visual motion was task relevant than when task irrelevant (Guerreiro et al., 2022). While this study suggested that overall excitability of the extrastriate cortical region MT can be top-down regulated, the present study demonstrates that the information send via feedback connections is precise enough to represent category specific visual information in V1.

Thus, on the one hand the existing feedback connections in CC individuals seem to be sufficient to encode typical features of a category. On the other hand, they seem to be too coarse to encode specific individual scenes as suggested by a lack of successful decoding of scene context in the occluded condition. This conclusion is supported by the connective field results.

Unlike in the SC group, the cortico-cortical spatial sampling extent in FF direction was not larger than in the FB direction during visual stimulation in the CC group. The absence of this typical convergent state during visual stimulation in CC individuals, which results from the absence of the typical state-dependent shift (de Best et al., 2020) from divergent processing during rest provides independent evidence for a lower tuning and thus precision of feedback connections. This is further supported by the significant association of CF sizes in FB, but not FF, direction with decoding accuracies.

### Coarser feedback processing results in impaired feedforward processing

Non-human animal studies have shown that RF size in V1 crucially depends on feedback connections from V2 (Nurminen et al., 2018). Since RF size defines the resolution of feedforward processing, e.g. visual acuity (Heitmann et al., 2023; Silva et al., 2021), a coarser FB connectivity must result in deficits in feedforward processing. This has impressively been demonstrated in a patient group with atrophies in posterior brain regions from which feedback connections to early visual cortex originates (de Best et al., 2020). These patients have intact retinas and feedforward activity to the primary visual cortex. Nevertheless, pRF sizes in the foveal region of V1 were larger and these patients typically report foveal crowding. Similarly, we had observed that pRF size of the fovea representing region of early visual cortex was larger in CC individuals and V1 pRF sizes correlated with visual acuity (Heitmann et al., 2023).

The present result of a larger spread of FB connectivity provides strong evidence that the larger pRF sizes of CC individuals in V1 were indeed due to impaired recurrent processing rather than aberrant feedforward input. In fact, the CF analysis strongly confirmed this idea by showing that under visual stimulation early visual cortex of CC individual was not able to switch into a convergence mode that would allow for the pooling of information downstream.

The results of the present study thus provide direct evidence from two methodological approaches for a disrupted orchestrating of feedback and feedforward processing as originally postulated as a consequence of deafness in cats (Kral et al., 2017; Yusuf et al., 2022) and as suggested by neurophysiological recordings in congenital cataract reversal individuals (Pitchaimuthu et al., 2021). Finally, anatomical studies in enucleated monkeys found a flattened visual cortical hierarchy due to aberrant feedback connectivity (Magrou et al., 2018) suggesting that the organization of the latter crucially depends on early experience. Recently, studies in rodents using visual deprivation (Dias et al., 2024) and selective rearing (Rajan et al., 2026) approaches have provided additional evidence for the experience dependence of feedback connectivity. Despite that feedforward and feedback connectivity were retinotopically aligned even in dark reared animals, the spatial overlap of input from the lateromedial visual area (which corresponds to V2 in primates) to V1 was larger. This observation is reminiscent of the larger CF sizes observed for V3/V2 to V1 (feedback connections) in the present study. Interestingly, Kowalewski et al. (2021) tested mice, either reared with standard visual input, dark reared or initially dark reared and exposed to visual input only as adults. The latter model approximates our model of congenital cataract reversal (although sight recovery was relatively earlier in CC individuals). The authors assessed natural scene discriminability in V1 which has been earlier demonstrated to arise from natural scene specific surround modulation (and thus, presumably feedback connections; Pecka et al., 2014). In an impressive correspondence to the results, we reported here in CC individuals, initially dark reared mice with consecutive 2-4 weeks of visual experience in adulthood featured lower scene discriminability in V1.

Individuals with reversed congenital cataracts were found to exhibit an altered ratio of alpha and gamma activity in EEG recording: Alpha and gamma activity have been considered to be interdependent, that is, alpha activity was postulated to indicate a top-down mechanism that regulates bottom-up activity which was postulated to show up as gamma activity (Popov et al., 2017; Wang et al., 2016). In fact, CC individuals’ resting state EEG was characterized by lower alpha oscillatory activity and higher gamma activity compared to normally sighted controls. This altered ratio was interpreted as impaired feedback processing that resulted in excessive feedforward activity (Ossandón et al., 2023; Pant et al., 2023).

In fact, the extensive work in mice in recent years has provided consistent evidence that visual input or patterned feedforward activity (Ibrahim et al., 2021) is crucial for the shaping of feedback activity. The neural networks resulting from this loop likely reflect the statistical structure of the visual input (see Kremers & Rose, 2024) and are thus particularly prepared to efficiently process the “expected” visual input (Berkes et al., 2011; Kowalewski et al., 2021).

### Visual acuity and higher visual functions rely on overlapping neural mechanisms

Currently, no evidence exists that the lower visual acuity characteristic for CC individuals arises at the retina. Instead, as discussed above, visual acuity deficits of the CC group more likely originate from neural circuit properties in early visual cortex. This idea is reminiscent of modelling work in neonates which suggested that their compared to adults’ lower visual acuity cannot fully be explained by the immature optics and retinae but rather must include immature neural circuits (Banks and Bennett, 1988).

The exploratory correlation analysis of the present study suggested that better visual acuity across groups was associated with larger convergence magnitudes during visual stimulation. Notably, the latter association was driven by both, larger average FF CF sizes and smaller average FB CF sizes. Thus, better visual acuity seems to be associated with both more extensive pooling downstream and more spatially precise upstream modulation. These results are in accord with the idea that RF size depends not only on feedforward but additionally on feedback processing (Nurminen et al., 2018), given that RF size has often been found to correlate with visual acuity (Heitmann et al., 2023; Silva et al., 2021). Here we hypothesize that the lower visual acuity of individuals with reversed congenital cataracts is due to both a lower feedforward convergence and a spatially less precise feedback modulation of early visual cortex. In many previous studies on higher visual functions in CC individuals, visual acuity differences compared to normally sighted controls have been considered as a confound: authors tried to control for the lower visual acuity of CC individuals by including visual acuity as a covariate (Ossandón et al., 2025), by blurring the images for normally sighted controls (Gilad-Gutnick et al., 2024; McKyton et al., 2015; Rajendran et al., 2020) or by including visual impaired controls (Röder et al., 2013). Here we postulate, that lower visual acuity and worse performance in higher visual functions such as face identity processing (Le Grand et al., 2001; Putzar et al., 2010) might be due to the same neural mechanisms.

Visual processing at all levels of the visual cortical hierarchy, depends on recurrent processing. Thus, the validity of blurring stimuli to estimate the role of visual acuity of CC individuals should be reconsidered. In the present study, visual acuity correlated with decoding accuracy of both scenes and categories for the non-occluded but not for the occluded ROIs. Thus, given the correlations of visual acuity with convergence magnitude, we postulate that lower visual acuity and lower decoding accuracies in CC participants might be due to a lower convergence magnitude, that is an altered ratio of FF and FB CF size in early visual cortex and thus might reflect the same neural mechanism. Notably, all CC individuals who behaviorally responded (6 out of 7), correctly identified the visual category context in the behavioral task, suggesting sufficient visual acuity to process the stimuli’s categories (i.e., highways, beaches, and buildings).

Removing effects of visual acuity and their underlying neural mechanisms left the correlation between decoding accuracies in the occluded ROI with the cortical magnification factor (CMF) significant in the CC group. As previously shown, CC individuals seem to particularly exhibit a coarser cortical representation of the foveal region compared to SC individuals, which was reflected by larger foveal pRF sizes and smaller CMF (Heitmann et al., 2023). Recently, the central-peripheral dichotomy theory (Zhaoping, 2019) postulated that the foveal region entertains a higher density of feedback connections than more peripheral regions. As a consequence, foveal input is particularly prevalent in ventral stream regions important for e.g. face processing (Finzi et al., 2021). This idea integrates reports in which information about peripherally presented objects was present in fovea-representing V1 (Bennett et al., 2025; Williams et al., 2008) and revives the idea of foveal V1 as a “cognitive active blackboard” (Mumford, 1992; Roelfsema & De Lange, 2016). We speculate that the disrupted ratio of smaller sampling in FF direction and (or due to) coarser feedback processing in CC individuals as demonstrated in the present study during visual stimulation might result in a coarser “foveal blackboard” affection all levels of visual processing.

### Limitations

Studying visual processing in congenital cataract reversal individuals constitutes a rare opportunity to gain knowledge about the role of early visual experience in human visual brain development. Research in this population directly links to the large body on non-human animal work on the effects of visual deprivation on visual brain development (Hubel & Wiesel, 1964; Levelt & Hübener, 2012).

CC individuals fulfilling the inclusion requirements (i.e., lack of visual pattern vision for at least 6 months after birth, adult age, no neurological condition, no high-field MRI contraindications and willingness to participate) are rare in Western countries (Röder et al., 2021), naturally limiting the sample size since the participants were recruited in Germany and scanned at the University of Maastricht (The Netherlands). While a greater sample size would be desirable, we are confident that our results add valuable insights. First, we were able to replicate previous findings of successful decoding of scene and category context in occluded and non-occluded ROIs (Morgan et al., 2019; Smith & Muckli, 2010). Second, we demonstrated a state-dependent shift in convergence magnitude in our group of SC individuals as has been reported previously (de Best et al., 2020). Third, our results were motivated by previous electrophysiological evidence suggesting impaired feedback processing in individual with reversed congenital cataracts (Bottari et al., 2016; Ossandón et al., 2025; Pant et al., 2023; Pitchaimuthu et al., 2021; Putzar et al., 2007).

Sight recovery individuals with a history of congenital visual deprivation typically present with nystagmus if the surgery was not performed within the first 8 weeks of congenital blindness (Lambert et al., 2006; Rogers et al., 1981). Since in the occlusion paradigm, the selection of vertices that did not receive any bottom-up visual information is critical, it is important to make sure that the receptive fields of these vertices did not “move” through stimulated parts of the presentation window during involuntary eye movements. By functionally defining the ROIs (see Methods), we applied conservative inclusion criteria to address this concern. In addition, excluding vertices falling into the “surround” region (see Figure 1B) from our ROIs minimized the influence of lateral connections. Importantly, the CF analysis corroborated and extended the results from the occlusion paradigm providing additional evidence for coarser feedback activity.

### Conclusion

Using two analysis approaches, task-based fMRI and CF analysis, we demonstrated a top-down modulation of early visual regions in CC individuals that was sufficient to encode different types of visual scenes (here categories), but not sufficient for encoding individual scenes. We hypothesize that lower convergence in downstream processing and diffuser feedback connectivity result in coarser processing in V1. We postulate that from this imbalance of ordered feedforward and spatially tuned feedback activity typical statistics of the visual environment are not properly coded resulting in typical impairments a less efficient processing across the visual processing hierarchy in individuals with reversed congenital cataracts.

## Data and Code Availability

Aggregated, pseudonymized data will be deposited at the University of Hamburg’s research data repository along with scripts to reproduce the analyses. Original data will be made available to external investigators upon reasonable request to the corresponding author through data transfer agreements approved by the stakeholders, under stipulations of applicable law including but not limited to the General Data Protection Regulation (GDPR; EU 2016/679).

## Author Contributions

- Conceptualization: B.R., C.H., M.L., R.G.
- Methodology: C.H., M.Z., R.V.H., B.R., R.G.
- Formal analysis and visualization: C.H., M.Z.
- Investigation: C.H., M.L., M.Z.
- Funding acquisition: B.R., R.G.
- Project administration: B.R., C.H., M.L., R.G.
- Supervision: B.R., M.Z.
- Writing – original draft: C.H., B.R.
- Writing – review & editing: R.G., C.H., M.Z., R.K.

## Acknowledgements

We thank Maria J. S. Guerreiro for her contributions in setting up the experiment, Suddha Sourav for the discussion on appropriate statistical analyses, Cordula Hölig for her support in preparing graphical illustrations and Tiago Mesquita for the exchange on connective field analysis.

