## Supplementary Material for "Early Vision Shapes Recurrent Processing in the Human Visual Cortex"

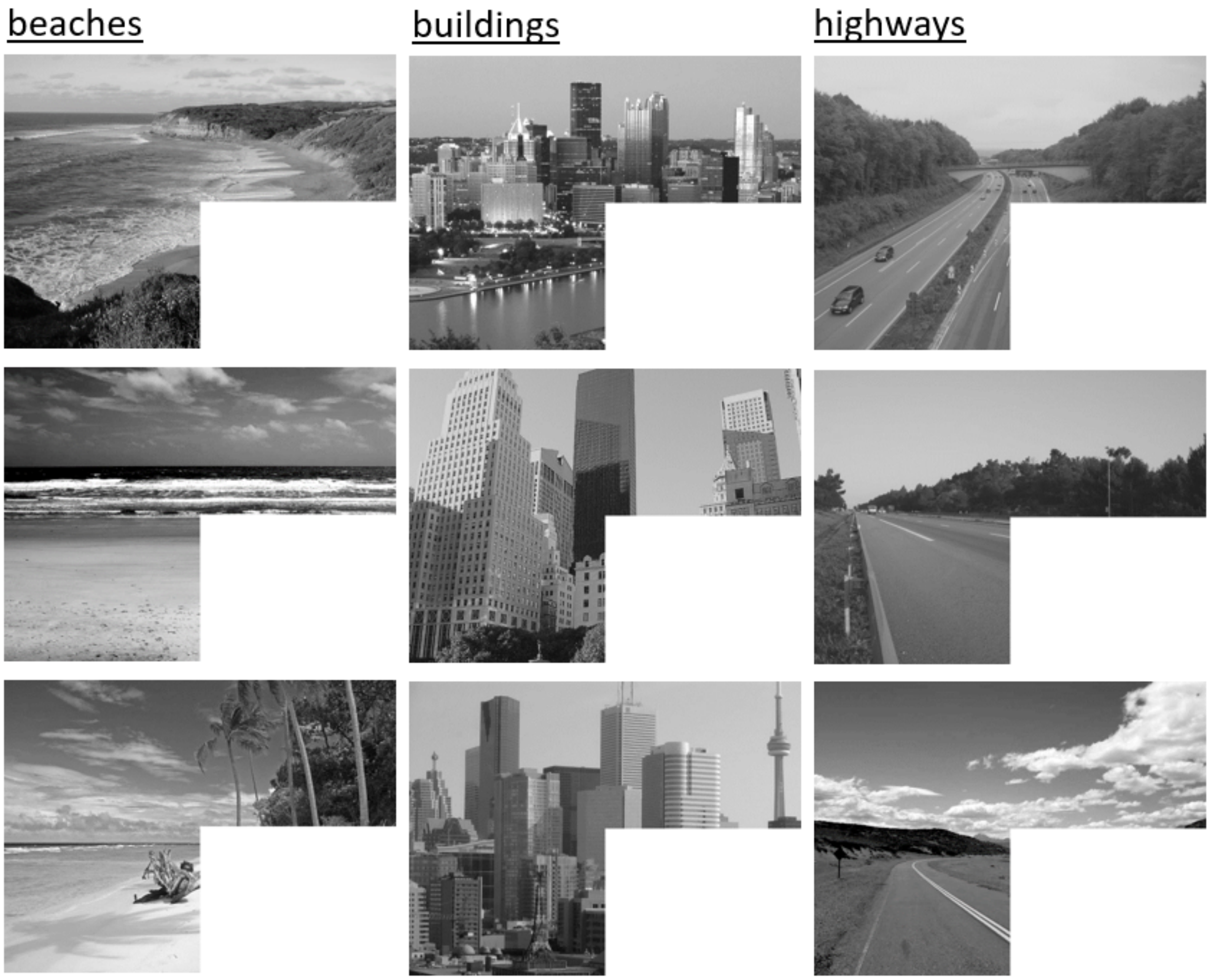


**Figure S1. Scene stimuli presented in this study.** An overview of the 9 grayscale scenes presented in this study. They belong to one of three categories of images: buildings, highways, and beaches.

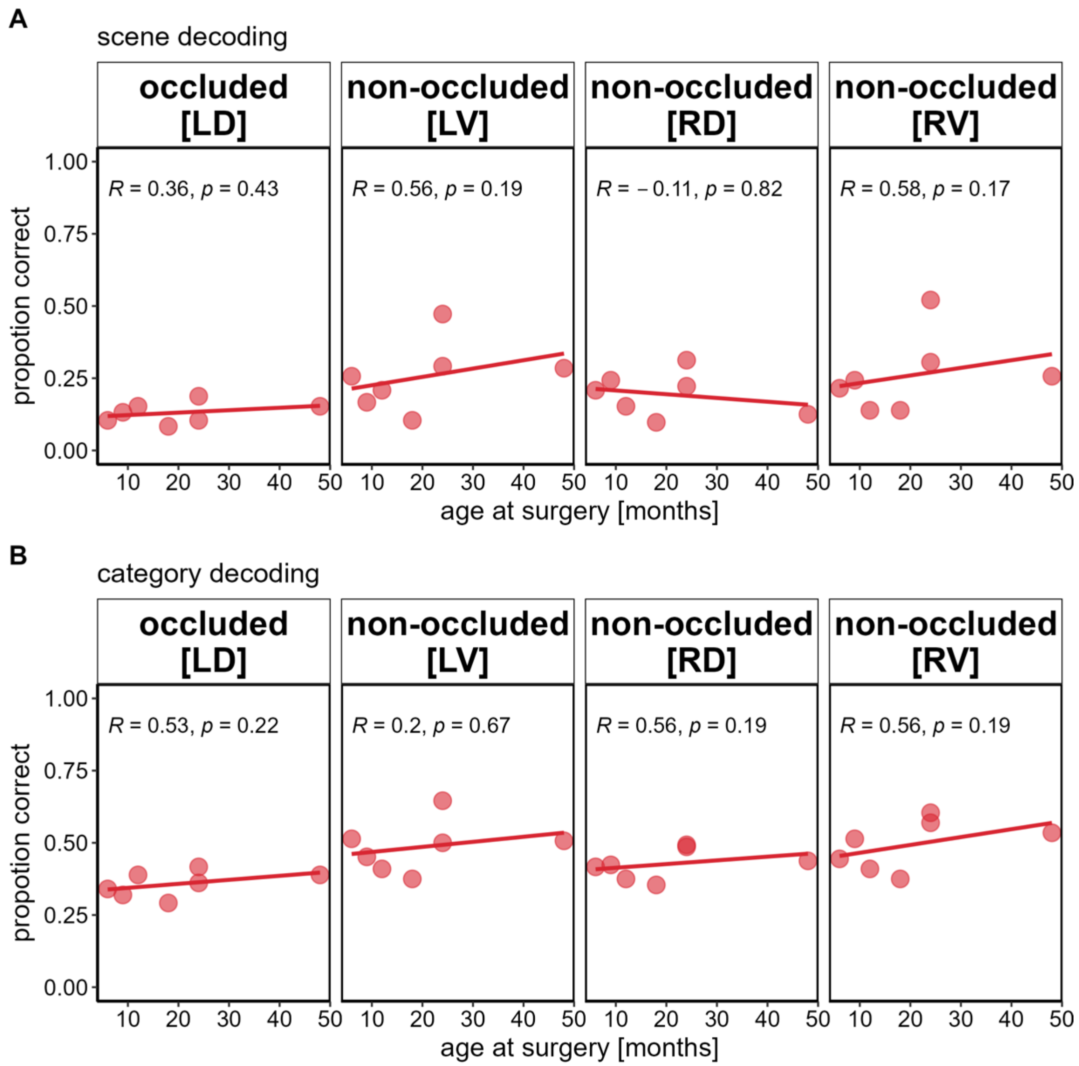


**Figure S2. Association of decoding accuracy and age at surgery separately per ROI.** Circles represent individual decoding accuracies in proportion of correct classifications for scenes (A) and categories (B). Spearman rank correlation coefficients and linear regression lines are inserted.

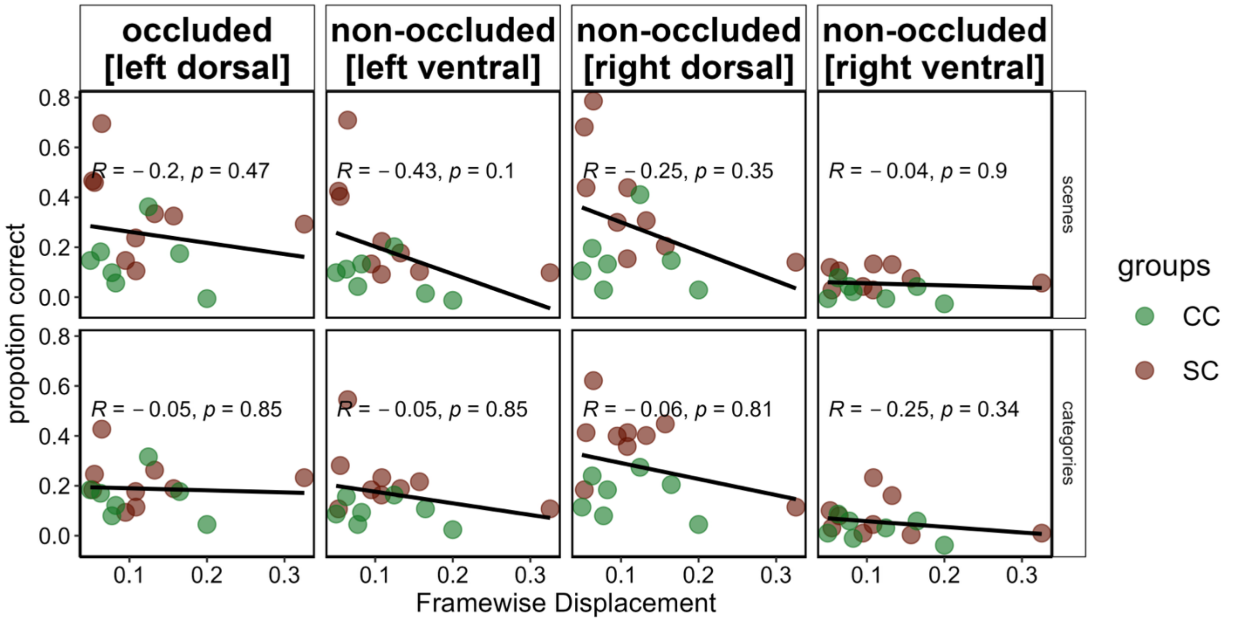


**Figure S3. Spearman's rank correlation coefficients between Framewise Displacement and decoding accuracy separately per ROI.** Circles represent individual decoding accuracies in proportion of correct classifications for scenes (upper row) or categories (bottom row) for target (left dorsal [LD]) and control (left ventral [LV] and right dorsal [RD]) regions. Spearman's rank correlation coefficients and linear regression lines are inserted across groups. The color of the circles indicate group (green = normally sighted control individual [SC], brown = congenital cataract reversal individual [CC]).

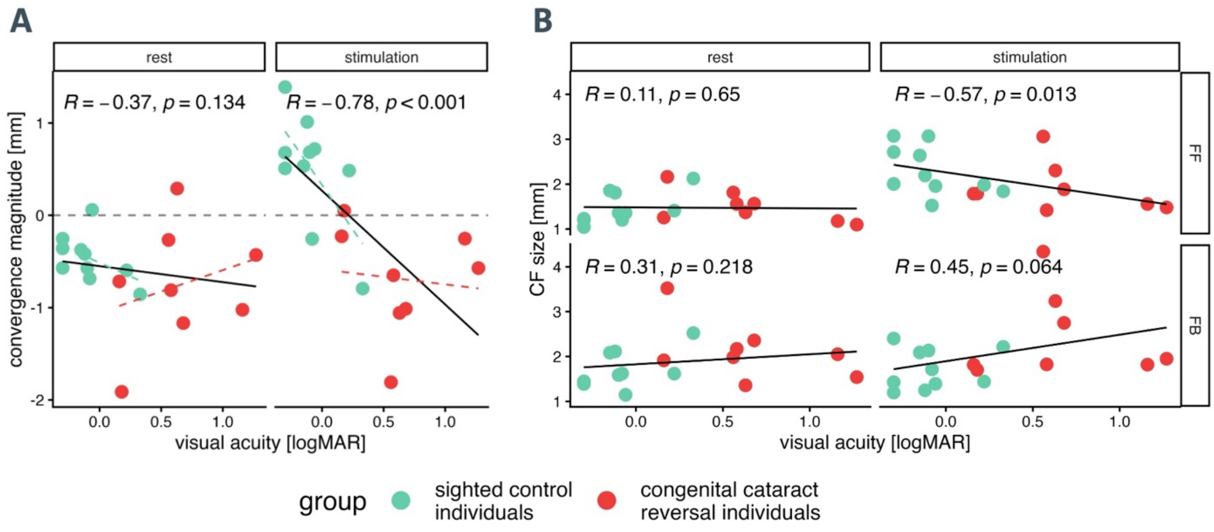


**Figure S4. Spearman correlation of visual acuity (in logMAR) and convergence magnitude (A), as well as average connective field sizes in feedforward and feedback processing direction per condition (B).** Circles represent the individual convergence magnitudes (A) or average CF sizes (B) in feedforward (FF) and feedback (FB) processing direction and visual acuity (in logMAR; lower values indicate better visual acuity) during rest (left panels) and stimulation (right panels). Spearman rank correlation coefficients and linear regression lines are displayed across groups (black).
